# Pathogenic Mechanisms Underlying Stargardt Macular Degeneration Linked to Mutations in the Transmembrane Domains of ABCA4

**DOI:** 10.1101/2020.08.28.272914

**Authors:** Fabian A. Garces, Jessica F. Scortecci, Robert S. Molday

## Abstract

ABCA4 is an ATP-binding cassette (ABC) transporter predominantly expressed in photoreceptors where it transports the substrate N-retinylidene-phosphatidylethanolamine across disc membranes thereby facilitating the clearance of retinal compounds from photoreceptor outer segments. Loss of function mutations in ABCA4 cause the accumulation of bisretinoids leading to Stargardt disease (STGD1) and other retinopathies. In this study, we examined the expression and functional properties of ABCA4 harboring disease-causing missense mutations in the two transmembrane domains (TMDs) of ABCA4. Our results indicate that these mutations lead to protein misfolding, loss in substrate binding, decreased ATPase activity or a combination of these properties. Additionally, we identified an arginine (R653) in transmembrane segment 2 of ABCA4 as a residue essential for substrate binding and substrate-stimulated ATPase activity. The expression and functional activity of the TMD variants correlate well with the severity of STGD1. Our studies provide a basis for developing and evaluating novel treatments for STGD1.

## Introduction

ABCA4 is a member of the A-subfamily of ATP-binding cassette (ABC) transporters primarily expressed in rod and cone photoreceptors (Allikmets, Singh et al. 1997, Illing, Molday et al. 1997, Molday, Rabin et al. 2000). Like other members of this subfamily, ABCA4 is organized as two nonidentical tandem halves with each half comprising a transmembrane domain (TMD), a nucleotide binding domain (NBD) and a large exocytoplasmic domains (ECD) (Bungert, Molday et al. 2001)(Fig 1). ABCA4 functions as a lipid importer flipping *N*-retinylidene-phosphatidylethanolamine (*N*-Ret-PE), the Schiff base adduct of retinal and phosphatidylethanolamine (PE), from the lumen to the cytoplasmic leaflet of photoreceptor outer segment disc membranes (Beharry, Zhong et al. 2004, Quazi, Lenevich et al. 2012). This transport activity insures that *all-trans* retinal (ATR) generated from photoexcitation and *11-cis* retinal not needed for the regeneration of rhodopsin and cone opsins are effectively cleared from photoreceptors by the visual cycle (Molday, Zhong et al. 2009, Quazi and Molday 2014).

**Figure 1.**
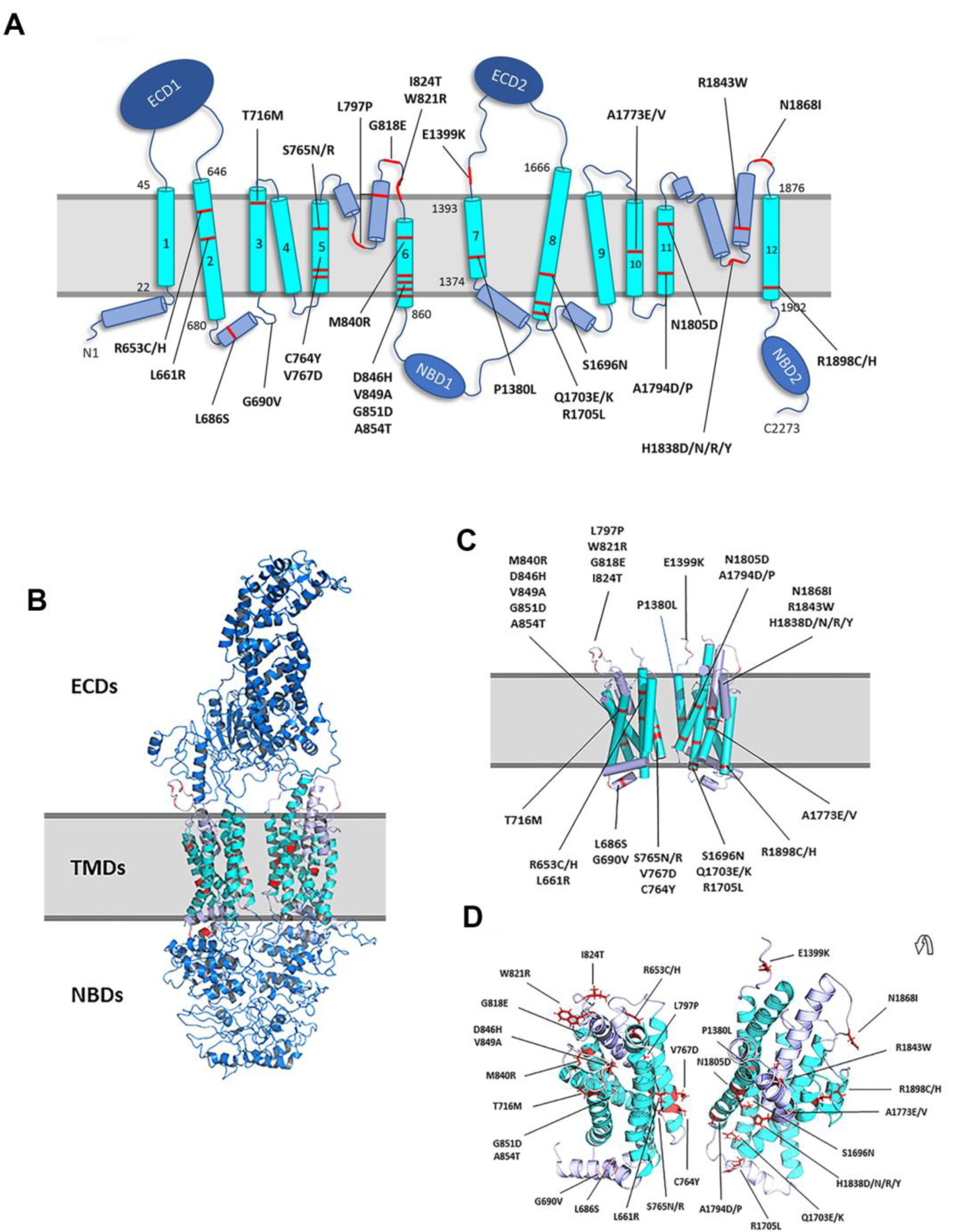
Location of TMD missense mutations in ABCA4 associated with STGD1. **A**. Topological model of ABCA4 showing the location of mutations in TMD1 and TMD2. **B**. (Left) ABCA4 structural model using the cryoEM structure of ABCA1 (5XYJ) as a template. The model highlights the ECDs (dark blue), TMDs (cyan for TM helices, and light blue for interconnecting loops/helices), and NBDs (dark blue). **C**. Depiction of the helical segments in TMD1 and TMD2 including the membrane spanning segments, cytoplasmic coupling helices, and exocytoplasmic V-shaped α-helical hairpin helices as cylinders together with the approximate location of disease-associated missense mutations. **D**. Top view of TMDs depicted as a ribbon cartoon together with the disease-associated mutations. ECD = Exocytoplasmic Domain; TMD = Transmembrane Domain; NBD = Nucleotide Binding Domain.

The importance of ABCA4 is highlighted by the finding that mutations in the gene encoding ABCA4 are responsible for autosomal recessive Stargardt disease (STGD1:MIM 248200) as well as recessive forms of cone-rod dystrophy and retinitis pigmentosa (Allikmets, Singh et al. 1997, Nasonkin, Illing et al. 1998, Klevering, Blankenagel et al. 2002, Charbel Issa, Barnard et al. 2013, Cornelis, Bax et al. 2017, Tanna, Strauss et al. 2017, Cremers, Lee et al. 2020). Stargardt disease, the most common inherited macular degeneration, is characterized by loss in central vision, progressive bilateral atrophy of the macula including the underlying retinal pigment epithelial (RPE) cells, impaired color vision, delayed dark adaptation, and accumulation of fluorescent yellow-white flecks around the macula and midretinal periphery. Over 1000 mutations in ABCA4 are now known to cause ABCA4-related diseases (https://databases.lovd.nl/shared/genes/ABCA4). These include missense mutations, frameshifts, truncations, small deletions, insertions, splice site mutations, and deep intronic mutations. Over 60% are missense mutations resulting in amino acid substitutions spread throughout the protein (Cornelis, Bax et al. 2017, Cremers, Lee et al. 2020).

A number of disease-associated mutations in the exocytoplasmic domains (ECDs), the nucleotide-binding domains (NBDs), and the C-terminal segment of ABCA4 have been previously characterized (Sun, Smallwood et al. 2000, Wiszniewski, Zaremba et al. 2005, Zhong, Molday et al. 2009, Quazi and Molday 2013, Zhang, Tsybovsky et al. 2015, Garces, Jiang et al. 2018). Many of these variants result in a significant reduction in cellular expression, mislocalization of ABCA4, and a loss in functional properties including substrate stimulated ATPase activity and ATP-dependent substrate transport activity. Loss of ABCA4 transport activity results in the formation and accumulation of bisretinoids including A2E in photoreceptors and RPE cells (Sparrow and Boulton 2005). These compounds accumulate as fluorescent liposfuscin deposits in STGD1 patients and animal models harboring *ABCA4* null alleles and loss of function mutations (Weng, Mata et al. 1999, Mata, Weng et al. 2000, Boyer, Higbee et al. 2012, Burke, Duncker et al. 2014, Zhang, Tsybovsky et al. 2015, Molday, Wahl et al. 2018, Makelainen, Godia et al. 2019). To date, however, the pathogenic mechanisms by which mutations in the TMDs of ABCA4 cause STGD1 have not been addressed in detail.

In this study we present comprehensive biochemical analyses of 38 STGD1 mutations localized to TMD1 and TMD2 of ABCA4 including the membrane spanning segments, a proposed coupling helix, and exocytoplasmic loops containing V-shaped hairpin helices (Fig 1). We show that most mutations in these regions predominantly affect the protein expression levels due to protein misfolding and significantly reduce *N*-Ret-PE binding and ATPase activity. The loss in activity can be used to predict the age of onset and severity of STGD1. As part of this study we have also identified an arginine at position 653 (R653) in transmembrane segment 2 (TM2) as a key residue required for the binding of *N*-Ret-PE to ABCA4.

## Results

### Transmembrane domains (TMDs) of ABCA4

Although earlier sequence analysis coupled with biochemical studies defined the topological organization of ABCA4 within membranes (Bungert, Molday et al. 2001), the exact location of the membrane spanning segments were not well-defined in this model. We have now used several programs that predict transmembrane segments including DAS-TMfilter, ExPASy TMpred, HMMTOP, MP Toppred, PredictProtein, and TMHMM (https://www.expasy.org/tools/). The results from these algorithms were pooled together in an effort to more definitively identify the membrane spanning segments (Figure 1 and Fig S1A-B). These assignments were further supported through analysis of the transmembrane segments defined in the cryo-electron structure of ABCA1 (Qian, Zhao et al. 2017). This member of the ABCA subfamily of ABC transporters shares over 50% sequence identity with ABCA4. This high degree of identity extends to the TMDs with TMD1 and TMD2 of ABCA4 sharing 51% and 60% identity with ABCA1, respectively (Fig S1A,B). On the basis of this high degree of sequence identity, we have generated a homology model of ABCA4 from the structure of ABCA1. The model shown Figure 1A-D is consistent with the membrane spanning segments predicted from transmembrane sequence algorithms. Not surprisingly, the ABCA4 model also predicts the location of intracellular transverse coupling helices (IH) and exoplasmic V-shaped α-helical hairpin helices (EH) as initially identified in ABCA1 (Qian, Zhao et al. 2017). The disease-associated mutations in the TMDs of ABCA4 analyzed in the present study are shown in Figure 1A,C,D and Fig S1A,B.

### Expression of TMD disease variants

The expression of 38 ABCA4 disease-associated variants in transfected HEK293T cells was quantified by western blotting after solubilization in the strong detergent SDS as a measure of total expression and after solubilization in the mild detergent CHAPS as a measure of more native-like protein. All ABCA4 variants with mutations in TMD1 and TMD2 expressed at similar levels when the cells solubilized with SDS were analyzed on western blots labeled for ABCA4 (Fig S2). However, significant differences in protein levels were observed for cells solubilized in CHAPS and subjected to high-speed centrifugation to remove aggregated protein prior to western blotting (Fig 2). Ten variants in TMD1 (L661R, L686S, G690V, S765N, S765R, V767D, L797P, M840R, D846H, G851D) solubilized below 40% WT ABCA4 levels; three (G818E, W821R, I824T) in the range of 40% to 52% WT ABCA4; and the remaining six (R653C, R653H, T716M, C764Y, V849A, A854T) at levels above 80% WT ABCA4 (Fig 2A., Table 1). In the case of TMD2, three variants solubilized at levels below 40% of WT ABCA4 (A1773E, A1773V, H1838R); seven variants between 40-80% of WT (P1380L, Q1703K, R1705L, A1794D, A1794P, H1838D, H1838Y); and nine variants (E1399K, S1696N, Q1703E, H1838N, N1805D, R1843W, N1868I, R1898C, R1898H) above 80% WT (Fig 2B. Table 2). These results suggest that many TMD variants consist of a significant fraction of highly misfolded, aggregated protein that fail to solubilize in CHAPS detergent resulting in the decreased levels of ABCA4 observed by western blotting.

**Table 1:** Summary of Biochemical Analysis of TMD1 Variants.

| Variant | Localization | Relative Solubilization<br>+/- SD | Basal ATPase Activity<br>+/- SD | N-Ret-PE induced ATPase Activity<br>+/- SD | N-ret-PE Binding +/- SD no ATP | N-Ret-PE Binding +/- SD with ATP | Class | Predicted Severity |
| --- | --- | --- | --- | --- | --- | --- | --- | --- |
| WT | Vesicles/ER | 100 | 100 | 224 +/- 22 | 100 | 15 +/- 7 | -- | Normal |
| R653C | Vesicles/ER | 102 +/- 9 | 113 +/- 21 | 134 +/- 17 | 17 +/- 12 | 15 +/- 10 | 1 | Severe |
| R653H | Vesicles/ER | 115 +/- 14 | 130 +/- 15 | 239 +/- 39 | 31 +/- 12 | 9 +/- 3 | 2 | Moderate |
| L661R | ER | 20 +/- 11 | 13 +/- 4 | 16 +/- 3 | 36 +/- 6 | 18 +/- 2 | 1 | Severe |
| L686S | ER | 27 +/- 8 | 18 +/- 9 | 30 +/- 16 | 16 +/- 8 | 11 +/- 7 | 1 | Severe |
| G690V | ER | 33 +/- 12 | 41 +/- 12 | 59 +/- 14 | 21 +/- 9 | 11 +/- 7 | 1 | Severe |
| T716M | Vesicles/ER | 98 +/- 16 | 94 +/- 4 | 228 +/- 13 | 59 +/- 7 | 9 +/- 8 | 3 | Mild |
| C764Y | Vesicles/ER | 100 +/- 12 | 133 +/- 5 | 270 +/- 39 | 68 +/- 6 | 16 +/- 7 | 3 | Mild |
| S765N | ER | 30 +/- 9 | 21 +/- 9 | 45 +/- 7 | 19 +/- 5 | 16 +/- 5 | 1 | Severe |
| S765R | ER | 29 +/- 11 | 18 +/- 6 | 29 +/- 4 | 13 +/- 4 | 18 +/- 3 | 1 | Severe |
| V767D | ER | 24 +/- 11 | 40 +/- 14 | 63 +/- 17 | 21 +/- 1 | 20 +/- 2 | 1 | Severe |
| L797P | ER | 25 +/- 5 | 31 +/- 5 | 37 +/- 2 | 11 +/- 5 | 12 +/- 5 | 1 | Severe |
| G818E | ER | 51 +/- 13 | 63 +/- 5 | 111 +/- 8 | 25 +/- 3 | 12 +/- 3 | 2 | Moderate /Severe |
| W821R | ER | 51 +/- 6 | 58 +/- 7 | 125 +/- 7 | 19 +/- 8 | 17 +/- 6 | 2 | Moderate /Severe |
| I824T | ER | 41 +/- 17 | 64 +/- 3 | 145 +/- 6 | 33 +/- 2 | 13 +/- 4 | 2 | Moderate /Severe |
| M840R | ER | 26 +/- 4 | 37 +/- 12 | 47 +/- 14 | 12 +/- 2 | 14 +/- 2 | 1 | Severe |
| D846H | ER | 30 +/- 9 | 42 +/- 22 | 50 +/- 18 | 13 +/- 12 | 14 +/- 7 | 1 | Severe |
| V849A | Vesicles/ER | 88 +/- 20 | 86 +/- 5 | 216 +/- 13 | 51 +/- 12 | 12 +/- 9 | 3 | Mild |
| G851D | ER | 27 +/- 11 | 31 +/- 11 | 46 +/- 12 | 13 +/- 8 | 12 +/- 10 | 1 | Severe |
| A854T | Vesicles/ER | 82 +/- 6 | 70 +/- 10 | 148 +/- 26 | 60 +/- 5 | 21 +/- 9 | 2 | Moderate |

Values are % of WT ABCA4 (100%)

**Table 2:** Summary of Biochemical Analysis of TMD2 Variants.

| Variant | Localization | Relative Solubilization<br>+/- SD | Basal ATPase Activity<br>+/- SD | N-ret-PE induced ATPase Activity<br>+/- SD | N-ret-PE Binding<br>+/- SD<br>no ATP | N-ret-PE Binding<br>+/- SD<br>with ATP | Class | Predicted Severity |
| --- | --- | --- | --- | --- | --- | --- | --- | --- |
| WT | Vesicles/ER | 100 | 100 | 222 +/- 16 | 100 | 16 +/- 7 | -- | Normal |
| P1380L | ER | 53 +/- 12 | 61 +/- 19 | 144 +/- 31 | 55 +/- 20 | 21 +/- 11 | 2 | Moderate |
| E1399K | Vesicles/ER | 102 +/- 15 | 94 +/- 11 | 202 +/- 34 | 117 +/- 13 | 31 +/- 14 | 3 | Mild |
| S1696N | Vesicles/ER | 96 +/- 17 | 120 +/- 8 | 255 +/- 7 | 116 +/- 16 | 27 +/- 13 | 3 | Mild |
| Q1703E | Vesicles/ER | 89 +/- 13 | 36 +/- 9 | 20 +/- 9 | 42 +/- 16 | 36 +/- 10 | 1 | Severe |
| Q1703K | ER | 52 +/- 11 | 17 +/- 2 | 12 +/- 1 | 37 +/- 6 | 35 +/- 20 | 1 | Severe |
| R1705L | ER | 40 +/- 13 | 76 +/- 5 | 113 +/- 9 | 54 +/- 12 | 21 +/- 9 | 2 | Moderate /Severe |
| A1773E | ER | 21 +/- 4 | 38 +/- 9 | 43 +/- 1 | 39 +/- 9 | 31 +/- 16 | 1 | Severe |
| A1773V | ER | 36 +/- 20 | 25 +/- 4 | 46 +/- 9 | 37 +/- 8 | 27 +/- 23 | 1 | Severe |
| A1794D | ER | 76 +/- 4 | 46 +/- 13 | 57 +/- 14 | 49 +/- 16 | 16 +/- 9 | 1 | Severe |
| A1794P | ER | 55 +/- 15 | 74 +/- 11 | 137 +/- 27 | 30 +/- 13 | 10 +/- 6 | 2 | Moderate /Severe |
| N1805D | Vesicles/ER | 82 +/- 18 | 74 +/- 15 | 154 +/- 19 | 44 +/- 11 | 18 +/- 3 | 2 | Moderate |
| H1838D | ER | 56 +/- 16 | 48 +/- 13 | 96 +/- 40 | 52 +/- 1 | 28 +/- 23 | 2 | Moderate /Severe |
| H1838N | Vesicles/ER | 107 +/- 14 | 60 +/- 6 | 132 +/- 13 | 48 +/- 2 | 33 +/- 10 | 2 | Moderate |
| H1838R | ER | 27 +/- 6 | 23 +/- 8 | 28 +/- 11 | 53 +/- 6 | 27 +/- 14 | 1 | Severe |
| H1838Y | ER | 48 +/- 11 | 48 +/- 18 | 75 +/- 20 | 34 +/- 8 | 16 +/- 3 | 1 | Severe |
| R1843W | Vesicles/ER | 99 +/- 8 | 68 +/- 3 | 131 +/- 14 | 84 +/- 11 | 27 +/- 14 | 3 | Mild/Moderate |
| N1868I | Vesicles/ER | 111 +/- 25 | 134 +/- 14 | 259 +/- 29 | 101 +/- 9 | 21 +/- 6 | 3 | Mild |
| R1898C | Vesicles/ER | 84 +/- 11 | 114 +/- 8 | 238 +/- 13 | 67 +/- 20 | 20 +/- 13 | 3 | Mild |
| R1898H | Vesicles/ER | 102 +/- 28 | 112 +/- 17 | 270 +/- 35 | 121 +/- 14 | 24 +/- 10 | 3 | Mild/Normal? |

Values are % of WT ABCA4 (100%)

**Figure 2.**
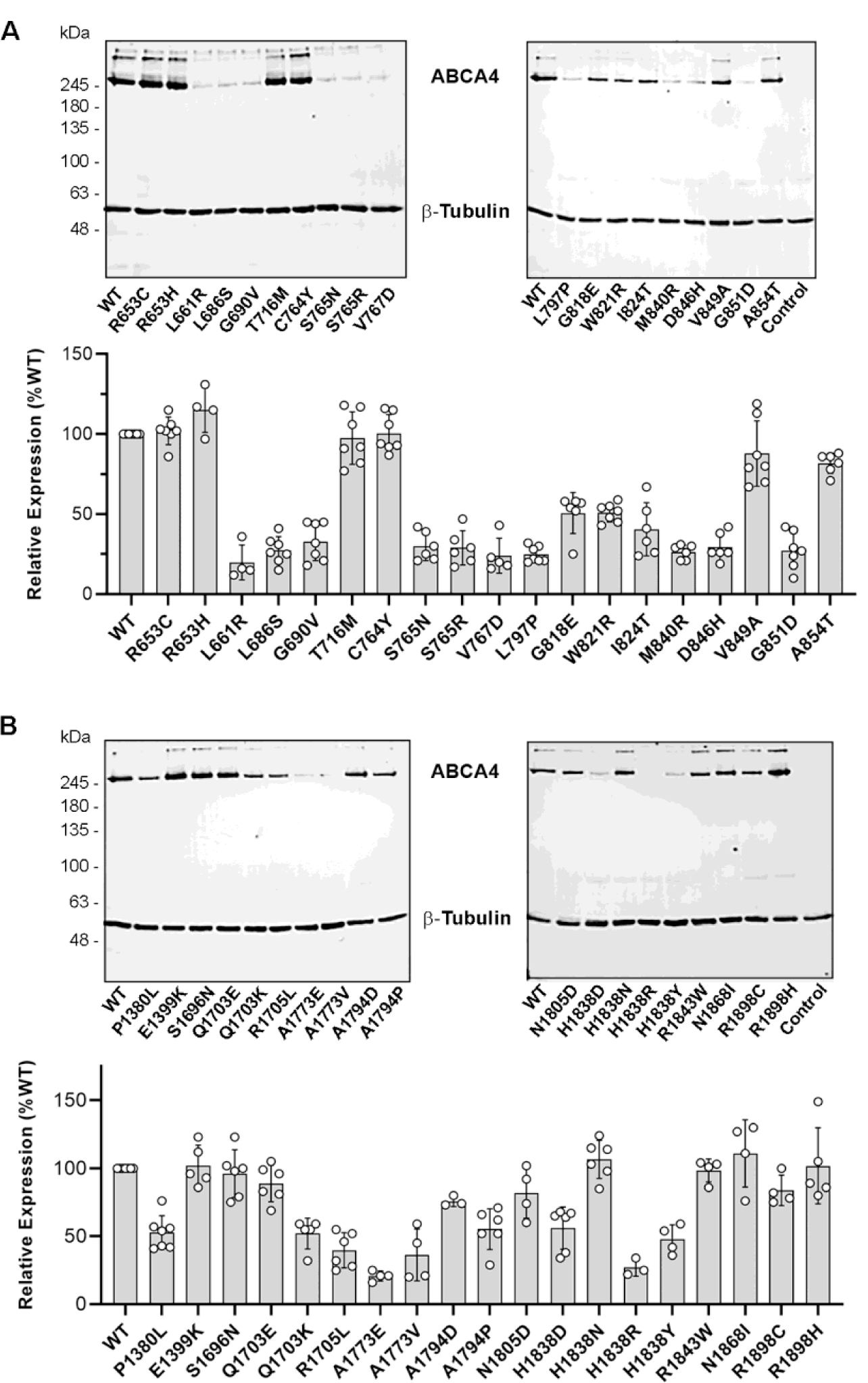
Solubilization of ABCA4 TMD variants. HEK293T cells expressing ABCA4 variants were solubilized in CHAPS. After removal of aggregated material by centrifugation, the cell supernatants were resolved by SDS gel electrophoresis and analyzed on Western blots labeled for ABCA4 and β-tubulin as a loading control. **A**. (Upper) Representative western blots of TMD1 variants. (Lower) Quantification of TMD1 variants relative to WT ABCA4. **B**. (Upper) Representative western blots of TMD2 variants. (Lower) Quantification of TMD2 variants relative to WT ABCA4. Control sample represent non-transfected HEK293T cells. Quantified profiles are the mean of relative expression levels ± SD for n≥4 where each point represents an independent experiment.

### Localization of ABCA Disease Variants in COS-7 cells

Previously, we showed that ABCA4 variants with low solubilization levels in CHAPS tend to be retained in the ER of transfected cells, whereas variants that express near WT levels accumulate in intracellular vesicle-like structures (Zhong, Molday et al. 2009, Garces, Jiang et al. 2018). Of the 38 variants examined in this study, 13 variants in TMD1 and 10 variants in TMD2 which solubilized in CHAPS at levels below 80% WT ABCA4 showed a reticular distribution in COS7 cells and co-localized with the ER marker calnexin, while TMD variants that solubilized above 80% WT levels displayed vesicular localization with some reticular labeling (Fig. 3A-C). These studies support the view that a significant fraction of the TMD variants that exhibit low solubilization in a mild detergent are misfolded and retained in the endoplasmic reticulum (ER) by the quality control system of the cell, while TMD variants showing high degree of solubilization by CHAPS can exit the ER and accumulate in vesicle-like structures similar to WT ABCA4.

**Figure 3.**
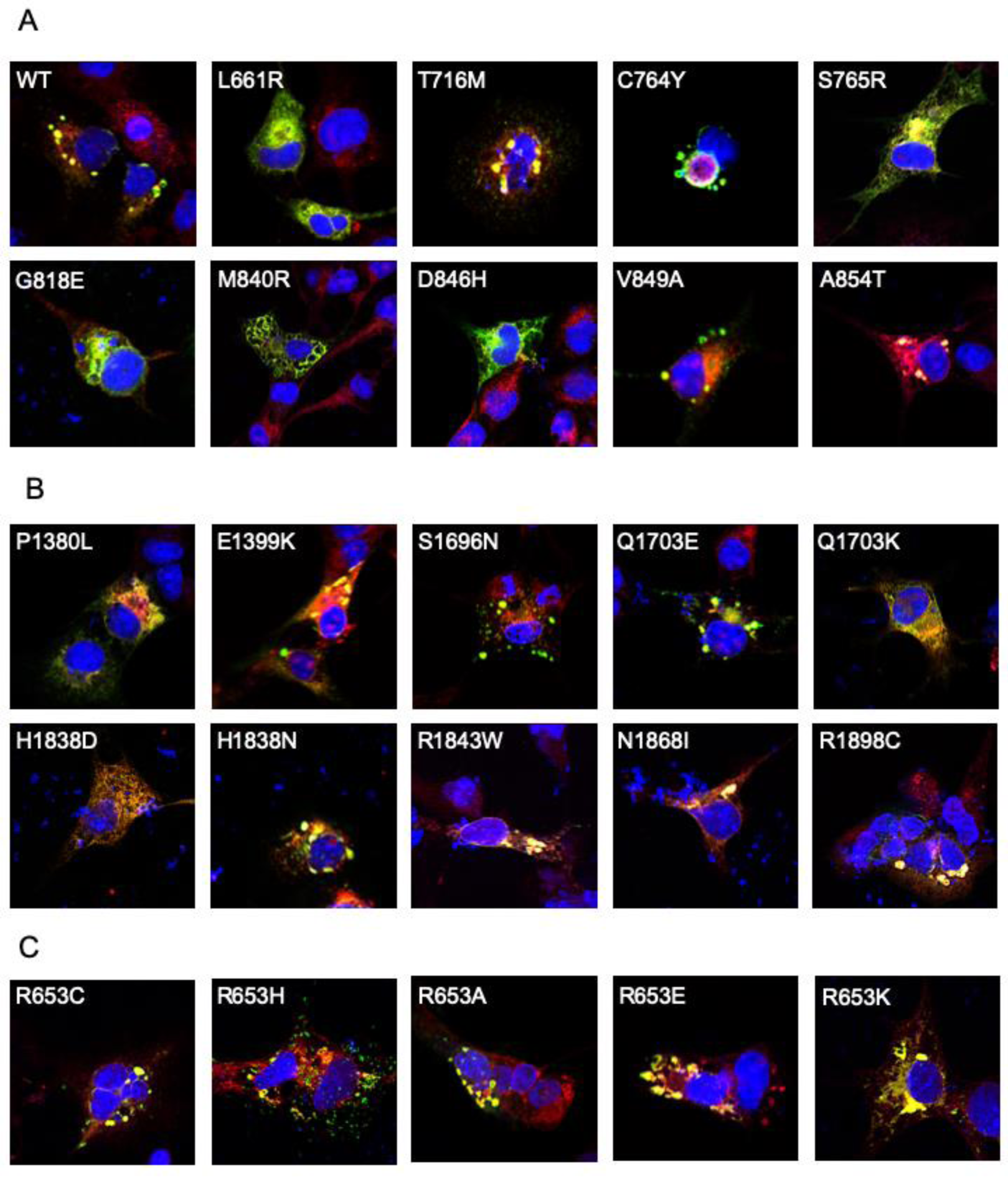
Cellular localization of TMD and R653 variants. Representative immunofluorescence micrographs of ABCA4 variants expressed in COS7 culture cells. The prevalence of vesicles is observed in cells expressing WT ABCA4 and variants with high levels of expression following CHAPS solubilization observed in Fig 2. Prevalence of reticular localization is observed in TMD variants with low solubilization. **A**. Representative TMD1 variants; **B**. Representative TMD2 Variants. **C**. R653 Variants. GREEN = rho1D4 antibody used to label ABCA4 variants containing the 1D4 tag; RED = anti-calnexin used as an ER marker; BLUE = DAPI (nucleus).

### Functional Analysis of TMD disease-associated variants: ATPase Assays

The ATPase activities and substrate binding properties of the ABCA4 variants were measured in order to determine the effect of TMD disease-associated mutations on the functional properties of ABCA4. For the ATPase assays, the ABCA4 variants expressed in HEK293T cells and solubilized in CHAPS detergent were purified on an immunoaffinity matrix. An aliquot of each variant was analyzed by SDS gel electrophoresis to ensure that that all variants had a similar degree of purity and concentration for use in the ATPase assays. All ABCA4 variants migrated as the major Coomassie blue stained band with an apparent molecular mass of 250 kDa supporting the view that all variants could be efficiently purified by immunoaffinity chromatography with a purity of over 90% (Fig. 4A).

**Figure 4.**
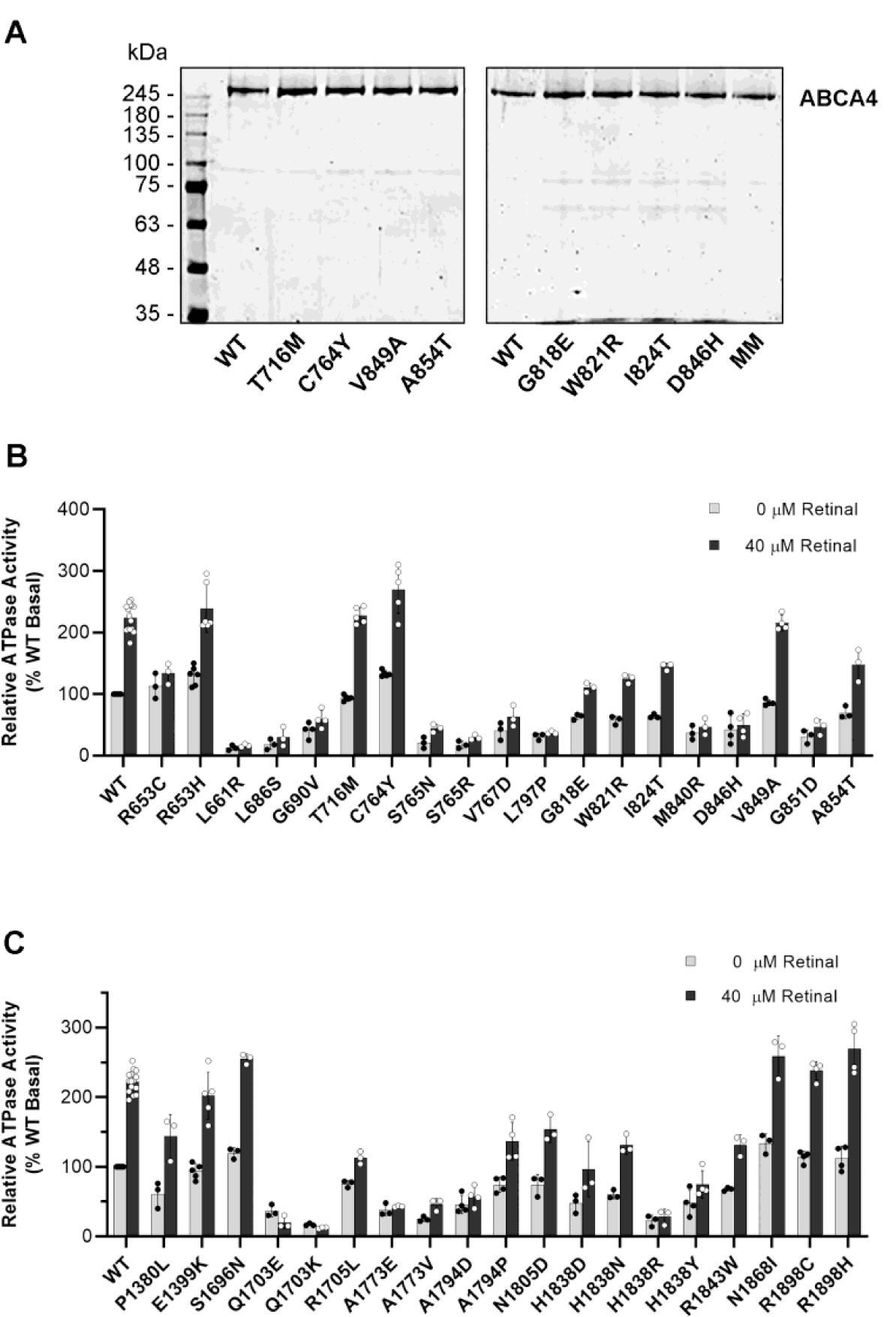
ATPase activity of ABCA4 TMD variants. TMD variants purified by immunoaffinity chromatography were analyzed for purity and protein concentration on SDS gels stained with Coomassie blue. **A**. Representative SDS gels of purified variants stained with Coomassie Blue. **B**. ATPase activity of the purified TMD1 variants in the absence and presence of 40 µM all-trans retinal; **C**. ATPase activity of the TMD2 variants. Data shows the mean activity ± SD for TMD1 variants n≥3 and for TMD2 variants n≥2. The ATPase-deficient variant (MM) in which the lysine residue in the Walker A motif of each NBD is replaced by a methionine residue was used as a control to correct for non-specific ATPase activity (see Fig 5)

The ATPase activity of detergent-solubilized WT ABCA4 and disease-associated variants was measured in the absence and presence of *all-trans* retinal (ATR). In the absence of ATR, but in the presence of phosphatidylethanolamine (PE), WT ABCA4 displays basal activity which largely reflects the energy-dependent flipping of the secondary substrate PE across membranes (Quazi and Molday 2013). Addition of 40 µM ATR in the presence of PE leads to the formation of the primary substrate *N*-Ret-PE and a two-fold increase in ATPase activity in agreement with previous reports (Sun, Molday et al. 1999, Ahn, Wong et al. 2000, Sun, Smallwood et al. 2000, Quazi, Lenevich et al. 2012, Garces, Jiang et al. 2018) (Fig 4B,C).

The ATPase activity of the TMD variants can be divided into three distinct groups. Group 1 includes variants that have basal ATPase activity below 50% of WT levels and show little or no *N*-Ret-PE induced stimulation in ATPase activity; Group 2 comprises variants with basal ATPase activity between 50-80% of WT levels and show modest *N*-Ret-PE stimulated ATPase activity; and Group 3 consists of variants that have both basal and *N*-Ret-PE stimulated ATPase activity comparable to WT. For TMD1, 10 variants (L661R, L686S, G690V, S765N, S765R, V767D, L797P, M840R, D846H, G851D) are classified in Group 1, 4 variants (G818E, W821R, I824T, A854T) belong to Group 2; and 4 variants (R653H, T716M, C764Y, V849A) are in Group 3 (Fig. 4B, Table 1). For TMD2 mutants, 7 variants (Q1703E, Q1703K, A1773E, A1773V, A1794D, H1838R, H1838Y) belong to Group 1; 7 variants (P1380L, R1705L, A1794P, N1805D, H1838D, H1838N, R1843W) to Group 2; and 5 variants (E1399K, S1696N, N1868I, R1898C, R1898H) to Group 3 (Fig. 4C, Table 2). The ATPase activities typically agreed with the degree of solubilization with some exceptions. In particular, the R653C variant expresses and displays basal ATPase activity similar to WT, but shows only minimal substrate-activated ATPase activity (Fig 4B).

To further define how various disease-associated mutations affect the ATPase activity of ABCA4, we measured the specific ATPase activity as a function of ATR concentration for selected Group 3 variants. For TMD1, T716M and V849A variants showed specific ATPase activity profiles similar to WT ABCA4, while C764Y variants had higher activity (Fig. 5A). The A854T variant in Group 2 which solubilizes in CHAPS at a WT level had a lower specific ATPase activity than WT (Fig. 5A). For the Group 3 variants in TMD2, four variants (E1399K, N1868I, R1898C and R1898H) had similar specific ATPase activities as WT while the S1696N mutant had higher specific activity (Fig. 5B). Also shown is the baseline activity measurements of the ATP-deficient MM double mutant (K969M/K1978M) (Ahn, Beharry et al. 2003) used to correct for any nonspecific luminescence signal generated in the ATPase assay.

**Figure 5.**
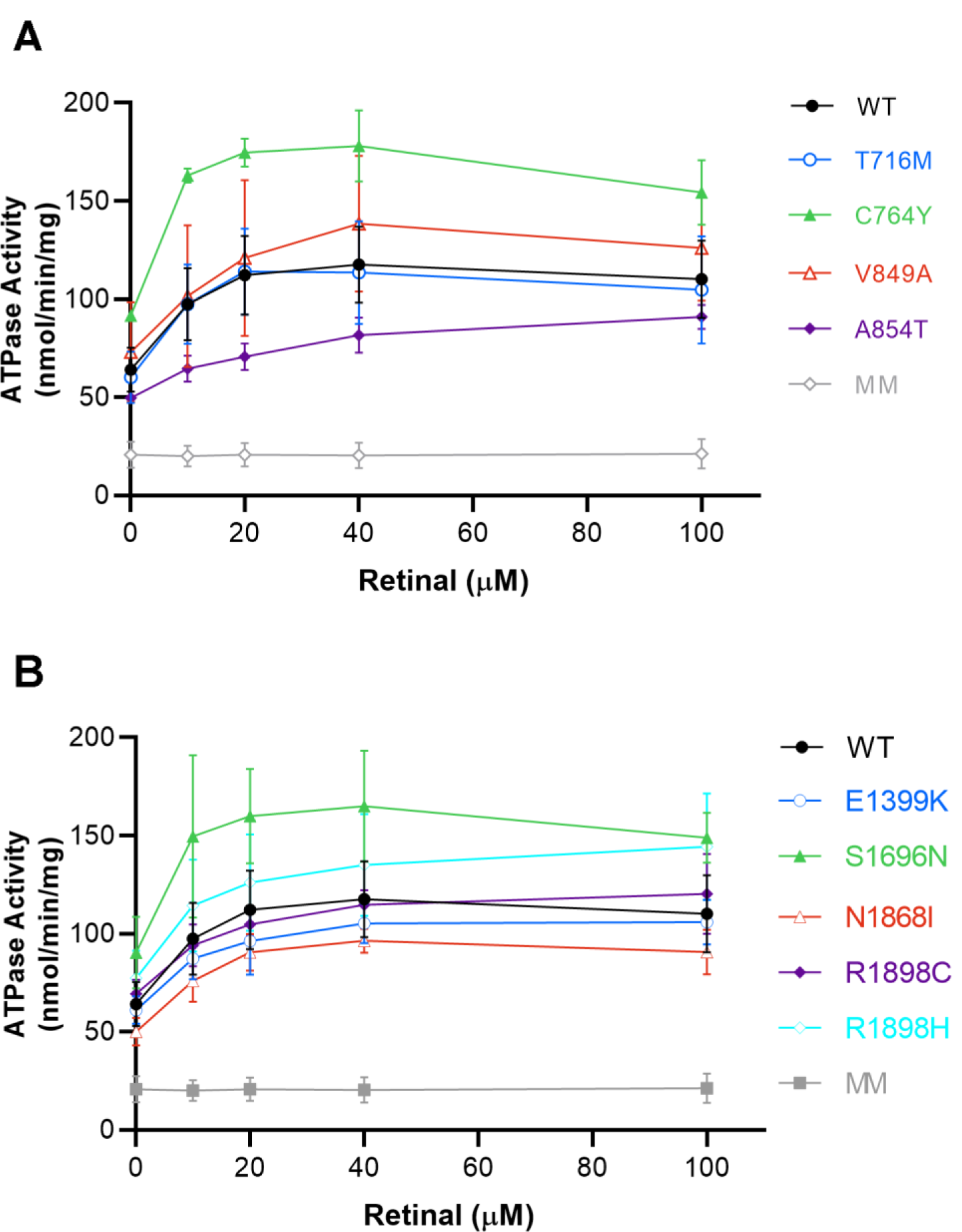
ATPase activity of selected TMD variants as a function of all-trans retinal concentration (Retinal). **A**. Profile of TMD1 variants. **B**. Profiles of TMD2 variants. Specific ATPase activity (nmol of ATP hydrolyzed per minute per milligram of ABCA4) was calculated for the TMD variants. Experiments were performed at concentrations similar to WT ABCA4 and show the mean activity ± SD for n≥2.

### Functional Analysis of disease-associated TMD variants: *N*-Ret-PE Binding Assays

The effect of TMD mutations on the binding of *N*-Ret-PE to ABCA4 was investigated using a solid phase binding protocol (Zhong, Molday et al. 2009, Garces, Jiang et al. 2018). ABCA4 immobilized on an immunoaffinity matrix was treated with ATR in the presence of PE to generate *N*-Ret-PE. After removal of excess substrate, the matrix was incubated in the absence or presence of 1 mM ATP and the amount of bound *N*-Ret-PE was determined. In the absence of ATP, *N*-Ret-PE tightly bound to WT ABCA4 as previously reported (Beharry, Zhong et al. 2004). The addition of ATP induced a conformational change in ABCA4 resulting in the efficient release of over 85% of the bound substrate.

The *N*-Ret-PE binding profiles for the TMD variants are shown in Figures 6A,B. Generally, the TMD variants could be separated into three main groups: One group bound *N*-Ret-PE at a level below 40% of WT levels, a second group bound *N*-Ret-PE in the range of 40-70% WT levels, and a third group bound *N*-Ret-PE above 70% WT levels. For TMD1, all 19 variants showed diminished *N*-Ret-PE binding. Fifteen variants (R653C, R653H, L661R, L686S, G690V, S765N, S765R, V767D, L797P, G818E, W821R, I824T, M840R, D846H, G851D) displaying very low substrate binding and little or no release in binding with ATP. These included the variants that had low basal and *N*-Ret-PE stimulated ATPase activity. Four variants (T716M, C764Y, V849A, A854T) bound *N*-Ret-PE in the range of 40-70% WT levels. For TMD2, however, the opposite was true with 15 of the 19 variants showing substrate binding above 40% WT ABCA4. Five of these variants (E1399K, S1696N, R1843W, N1868I, R1898H) displayed *N*-Ret-PE-binding and release similar or in some cases greater than WT ABCA4 (Fig. 6B, Table 2).

**Figure 6.**
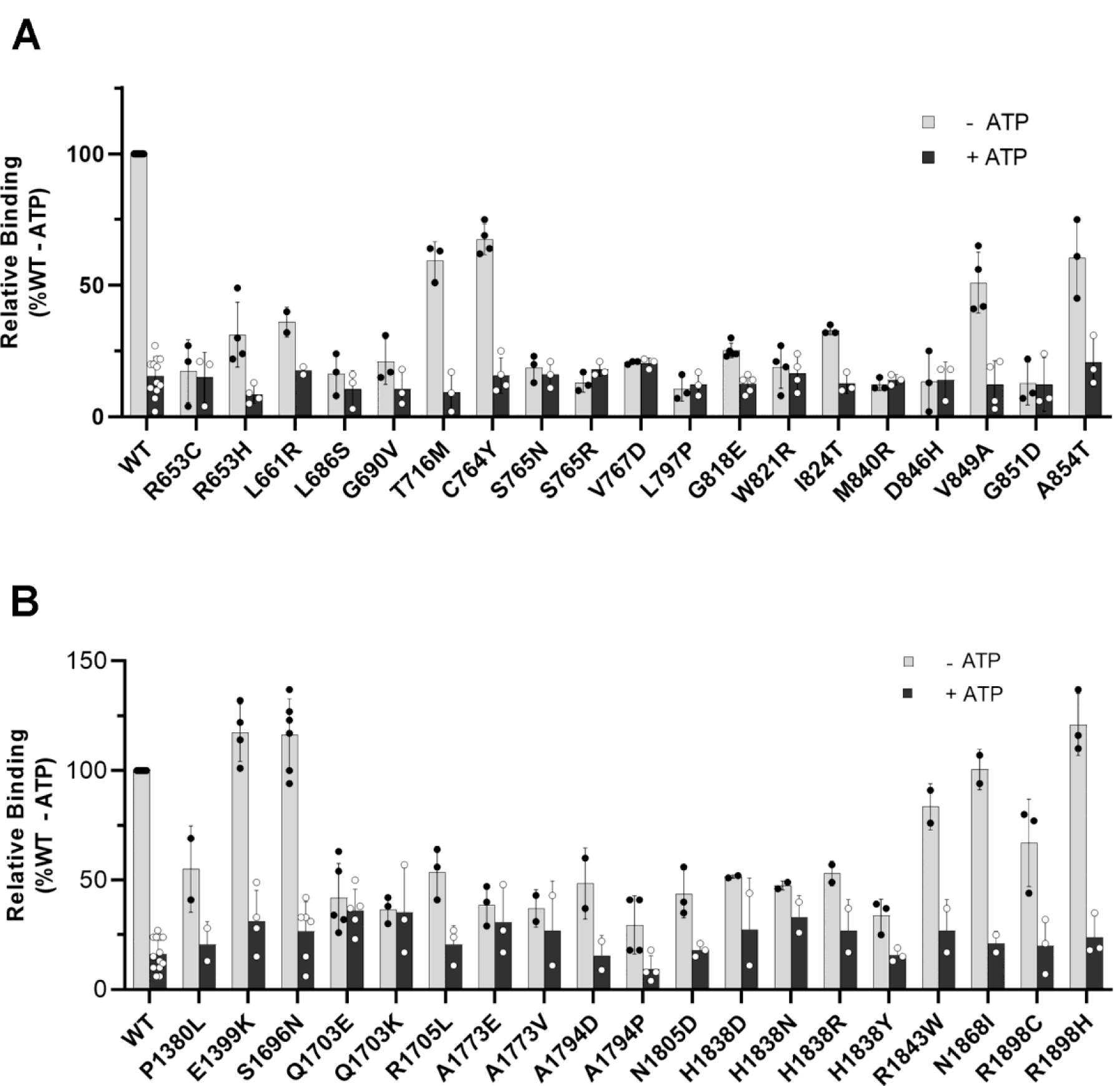
Binding of *N*-Ret-PE to TMD variants. ABCA4 TMD variants immobilized on an immunoaffinity matrix and containing bound *N*-Ret-PE were treated in the absence (grey) or presence (black) of 1mM ATP. **A**. *N*-Ret-PE binding profiles for TMD1 variants shown as a mean ± SD for n≥3 and **B**. TMD2 variants shown as a mean ± SD for n≥2.

Substrate binding curves were constructed to further investigate the apparent affinity of selected variants for *N*-Ret-PE. For the TMD1 variants T716M, C764Y, V849A, and A854T, the apparent dissociation constant K_d_ was 4 to 7 times higher than for WT ABCA4, indicating that these variants had significantly lower affinity for its substrate than WT ABCA4 (Fig. 7A). For the TMD2 variants, there was a wide range in K_d_ values. The apparent K_d_ for the E1399K and S1696N variants was 2 and 3 times higher than for WT ABCA4, while the K_d_ for the N1868I, R1898C, and R1843W variants were 12, 14 and 26 times higher, respectively (Fig. 7B). These results indicate that the affinity of the TMD variants for *N*-Ret-PE varies widely with many showing only a modest decrease in affinity of 2 to 5 fold (K_d_ 2.8-6.3 µM) and others showing a significantly larger decrease in affinity (K_d_ 8.6-31.1 µM). These results also emphasize that both TMDs contribute to the efficient binding and release of *N*-Ret-PE.

**Figure 7.**
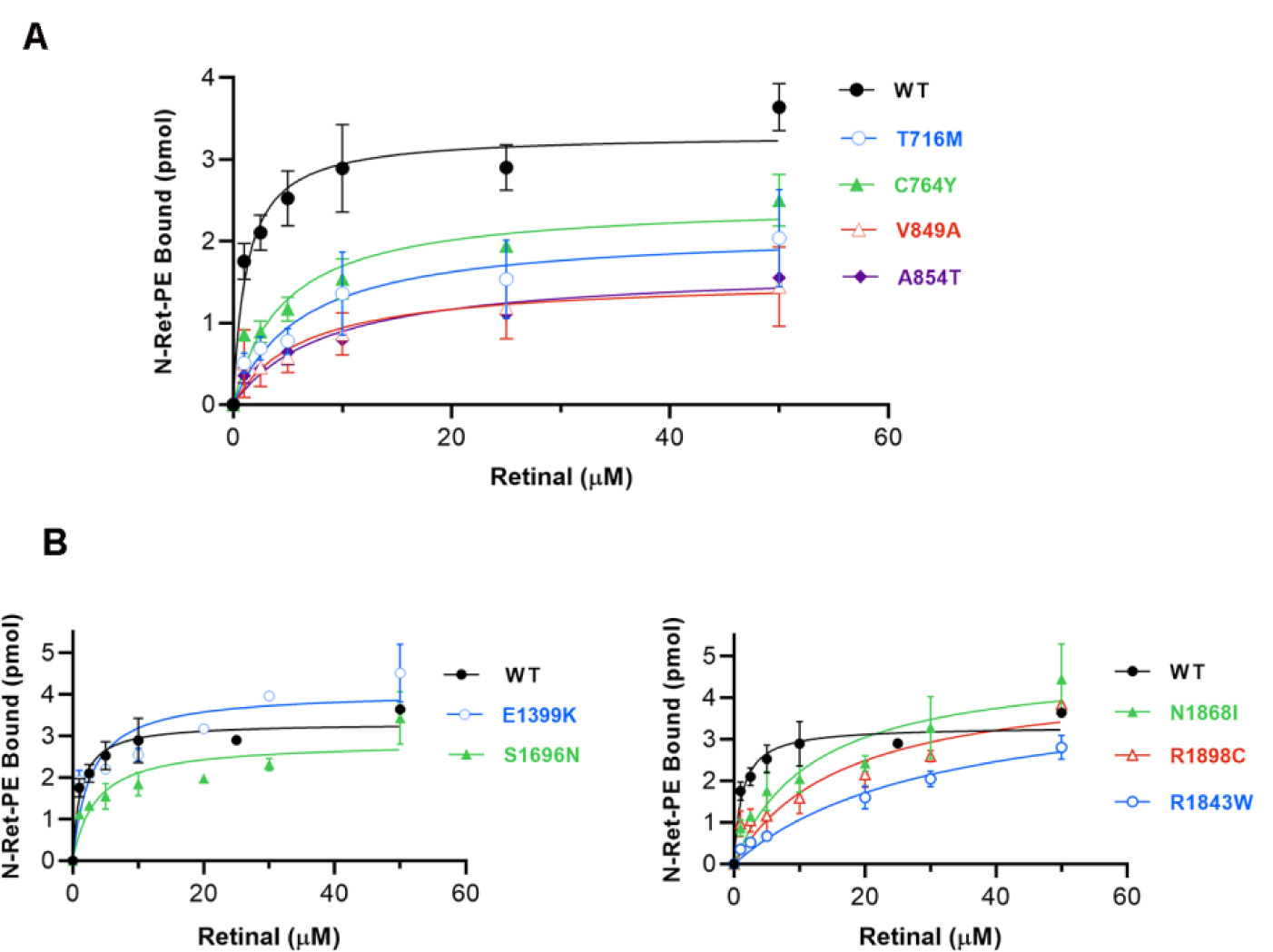
Binding of *N*-Ret-PE to selected TMD variants as a function of all-*trans* retinal concentration. ABCA4 TMD variants immobilized on an immunoaffinity matrix were treated with various concentrations of radiolabeled all-*trans* retinal. The amount of bound *N*-Ret-PE was determined after removal of excess retinal. **A**. Best fit binding curves for TMD1 variants (n≥2) for the following apparent mean K_d_ ± SD; WT K_d_ = 1.2 ± 0.2 µM; T716M K_d_ = 6.3 ±0.7 µM; C764Y K_d_ = 4.6 ±0.4 µM; V849A K_d_ = 6.4 ±0.9 µM; A854T K_d_ = 8.6 ±0.9 µM. **B**. Best fit binding curves for TMD2 variants for n≥2; K_d_ = E1399K K_d_ = 2.8 ±0.3 µM; S1696N K_d_ = 3.5 ± 0.3 µM; R1843W K_d_ =31 ± 3.5 µM; N1868I K_d_ = 12.0 ± 0.8 µM; R1898C K_d_ = 16.0 ± 1.7 µM.

### Biochemical Characterization of ABCA4 R653 Variants

The R653C and R653H disease-linked variants displayed distinct biochemical properties when compared to other TMD variants. Both variants expressed and solubilized in mild detergent at WT levels (Fig. 8A), displayed a WT-like vesicular expression pattern in transfected COS-7 cells (Fig. 3C), and retained a basal ATPase activity similar to WT ABCA4 (Fig 8B). However, the basal ATPase activity of the R653C variant was only marginally activated by *N*-ret-PE (19% increase for the R653C variant compared to 225% increase for WT ABCA4) as shown in Fig 8B and this variant was highly deficient in *N*-Ret-PE binding (Fig 8C). In contrast, the ATPase activity of the R653H variant was stimulated by *N*-Ret-PE, and this variant displayed significant substrate binding in the absence of ATP although at a reduced level relative to WT ABCA4 (Fig 8C). These properties prompted us to further explore the effect of other amino acid substitutions at position 653 of ABCA4 with the objective to determine if a positively charged residue at this position is required for substrate binding and ATPase activation.

**Figure 8.**
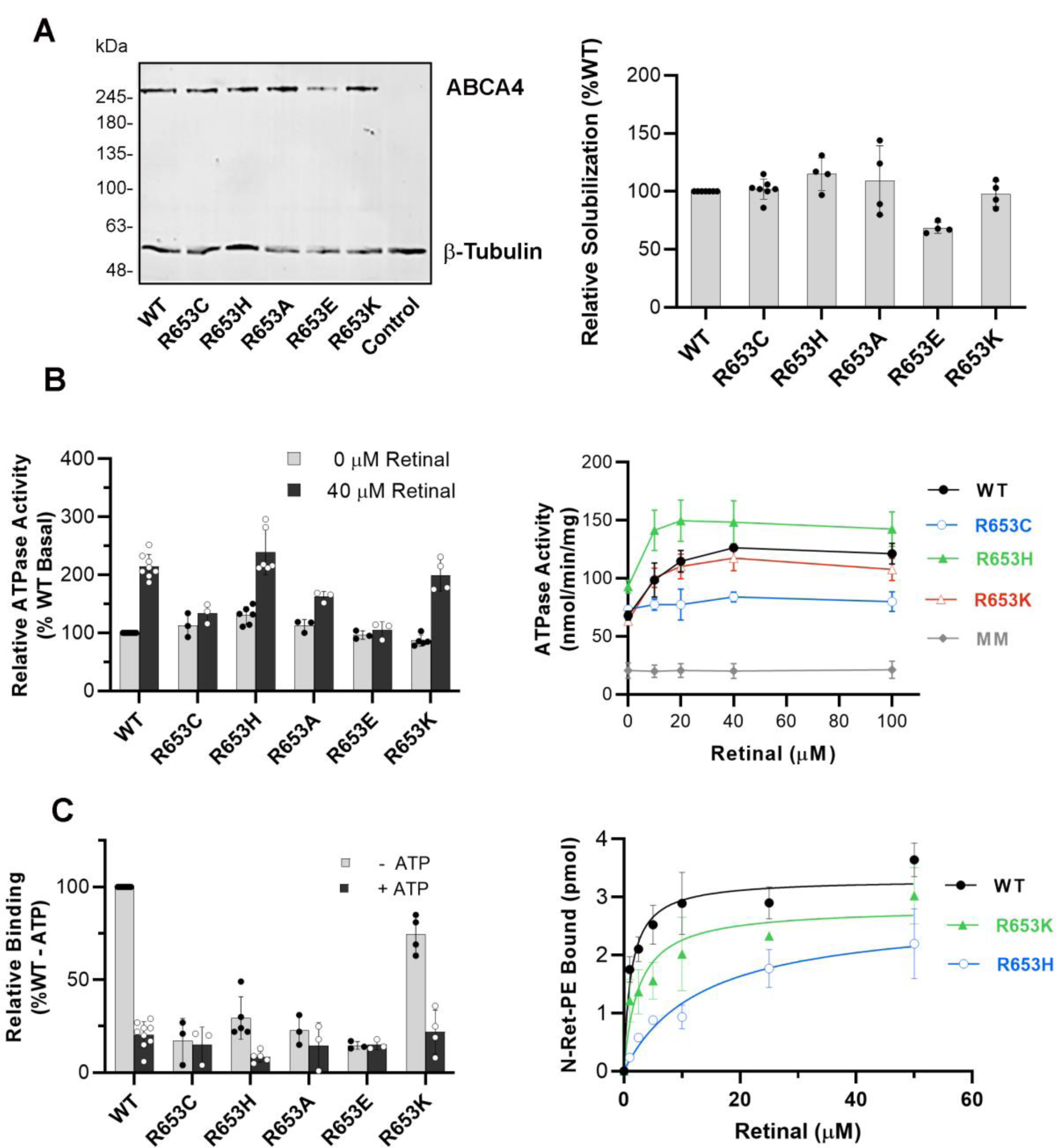
Biochemical analysis of R653 Variants. **A**. (Left) Representative western blots showing the CHAPS solubilization level of the R653 variants. (Right) Bar plot indicates the average relative expression of R653 variants ± SD (n≥2). **B**. (Left) Relative ATPase activity of R653 variants in the absence (grey) or presence (black) of 40 µM all-*trans*-retinal (n≥3). P-values with and without retinal: R653C P=0.021; R653H P=0.0005; R653A P=0.0005; R653E P value =0.426; R653K P=0.0013. (Right) The specific ATPase activity was measured for R653C/H/K variants as a function of all-*trans* retinal concentration (n≥2). The MM ATPase-deficient variant is also shown. **C**. (Left) Relative binding to *N*-Ret-PE in the absence (grey) or presence (black) of 1 mM ATP. P-values with and without ATP: R653C P value =0.422; R653H 0.0087; R653A P= 0.289; R653E P=0.833; R653K P = 0.0043. (Right) Best fit retinal binding curves for the variants (n≥2) with the following apparent mean K_d_ ±SD; WT K_d_ = 1.2 ± 0.2 µM; R653K K_d_ = 2.5 ± 1 µM and R653K K_d_ = 13.3 ± 2.5 µM

For these studies, three additional mutants containing a neutral (R653A), a negatively-charged (R653E), and a positively-charged (R653K) residue were constructed and their properties were compared to the R653C and R653H mutants. All these variants solubilized at or close to WT levels and formed vesicle-like structures in transfected COS-7 cells, indicative of proper protein folding (Fig. 3C and Fig. 4B,8A). They also retained basal ATPase activity levels similar to WT ABCA4 levels (Fig. 8B). However, only the variants with positively charged residues (R653H and R653K) displayed a significant *N*-Ret-PE induced increase in ATPase activity with the R653H variant showing statistically higher basal and substrate stimulated activity than even WT ABCA4 (Fig. 8B). The R653A variant, like the R653C variant, showed only a small although statistically significant increase in *N*-Ret-PE stimulated ATPase activity, and the R653E variant showed no increase (Fig 8B). *N*-Ret-PE binding properties of R653C, R653A, and R653E were drastically decreased, but only moderately reduced for R653K (Fig. 8C). The substrate binding affinity of these variants was also measured. The apparent K_d_ of R653K was only 2-fold greater than WT ABCA4 (K_d_ 2.5 µM for R653K compared to 1.2 µM for WT ABCA4) as shown in Fig. 8C. However, the R653H variant had a significantly lower affinity with a K_d_ of 13.3 µM. Taken together, these results indicate that a positively charged residue at position 653 is important for significant and stable *N*-Ret-PE binding and ATPase activation.

## Discussion

In this study, we analyzed 38 disease-causing missense mutations localized within the two TMDs of ABCA4 in order to define the pathogenic mechanisms responsible for STGD1. The majority of the mutations were amino acid replacements within the membrane spanning segments of TMD1 and TMD2 (Fig 1). Two mutations, however, were present in the cytoplasmic loop containing the intracellular transverse helix IH2 connecting TM2 and TM3 and eight mutations were situated in either of the two extracellular loops containing the V-shaped helices EH1 and EH2 between TM 5 and TM6 and EH3 and EH4 between TM11 and TM12 based on our homology model generated from the cryo-EM structure of ABCA1 (Fig 1 A-D). Our results indicate that most of the disease-causing TMD mutations lead to protein misfolding, decreased substrate binding, diminished basal and substrate-activated ATPase activity or a combination of these properties.

The residual activity of disease-associated ABCA4 variants and their level of expression are important factors in understanding protein structure-function relationships and pathogenic mechanisms underlying STGD1. The function of the ABCA4 variants is best determined by measuring the ATP-dependent flipping of *N*-Ret-PE across membranes, an activity required for the removal of toxic retinal compounds from photoreceptors (Quazi, Lenevich et al. 2012, Molday 2015). However, *N*-Ret-PE transport assays are difficult to carryout particularly when there is a need to evaluate large numbers of variants. Instead, we have used the activation of the basal ATPase activity by *N*-Ret-PE as a measure of ABCA4 function since this assay is straightforward to carryout, sensitive, and mirrors the ATP-dependent substrate transport activity of ABCA4 (Sun, Smallwood et al. 2000, Quazi, Lenevich et al. 2012, Quazi and Molday 2014). The degree to which a mild detergent solubilizes ABCA4 variants from membranes of transfected cells was used as a measure of the expression levels of the variants. The expression level of one disease-associated variant N965S in culture cells has been shown to agree with expression levels of ABCA4 in mice homozygous for this variant (Molday, Wahl et al. 2018) supporting the use of culture cells to measure mutant ABCA4 expression.

On the basis of ATPase activity measurements, substrate binding properties, and expression levels, the TMD variants can be separated into three major classes (Table 1 and 2). Class 1 consists of ABCA4 variants that typically show highly diminished expression and basal ATPase activities below 50% of the WT ABCA4 level with little if any *N*-Ret-PE stimulation of ATPase activity or *N*-Ret-PE binding and release by ATP. These loss-of-function variants are predicted to confer an early disease onset, typically within the first decade of life and a severe progressive visual impairment in individuals homozygous for these variants or compound heterozygous with another severe loss-of-function variant. Class 2 includes variants that have reduced expression and basal ATPase activities in the range of 50-80% WT ABCA4, but show modest *N*-Ret-PE binding and stimulation of basal ATPase activity. These variants are predicted to convey a more moderate disease phenotype with an age of onset in the second to fourth decade of life. Class 3 consists of variants with significant expression levels and basal ATPase activity typically greater than 80% WT levels and robust *N*-Ret-PE stimulated ATPase activity. These variants are predicted to impart a late disease onset and mild visual impairment. When paired with another mild variant these Class 3 variants may or may not confer a disease phenotype depending on the combined activity of these variants. In some instances, these variants may be considered as hypomorphic variants.

Of the 38 TMD variants studied here, 18 ABCA4 variants (11 in TMD1 and 7 in TMD2) comprise Class 1 (Table 1,2). In almost all cases, the low basal and substrate stimulated ATPase activity of these variants coincides with a loss in ATP-dependent *N*-Ret-PE binding and a low level of expression after mild detergent solubilization. These properties are consistent with these mutations causing extensive protein misfolding. This is further supported by the finding that these variants are retained in the ER of transfected cells by the quality control system of the cell as visualized by immunofluorescence microscopy. Indeed, most of these missense mutations involve amino acid substitutions which alter the charge or polarity of the side chains. Such a change can adversely affect the folding and packing of the transmembrane helices, the interface between the side chains and the hydrophobic lipid bilayer, or the conformation of loops connecting the membrane spanning segments. Interestingly, two disease-associated mutations (L686S and G690V) in Class 1 reside in the cytoplasmic loop between TM2 and TM3 containing a proposed intracellular coupling helix (IH2) which likely coordinates ATP binding and hydrolysis in the NBDs with transport of substrate through the TMDs (Fig 9A). Glycine at position 690 is conserved in ABCA4 from other vertebrates as well as ABCA1 and ABCA7 (Fig S1A). Leucine at position 686 has a nonpolar methionine at this position in ABCA1 and ABCA7 as well as in some ABCA4 orthologues supporting the importance of an amino acid with a hydrophobic side chain within this structural motif. One mutation (L797P) in Class 1 and three (G818E, I824T, W821R) in Class 2 are situated in highly conserved regions within the exocytoplasmic loop between TM5 and TM6, and another five (H1838D/N/R/Y and R1843W) in Group 1 and 2, reside in a conserved region within the exocytoplasmic loop between TM11 and TM12 (Fig S1,S2). These two loops contain the V-shaped α-helical hairpin helices as initially reported for the structure of ABCA1 (Qian, Zhao et al. 2017) and found in the homology model of ABCA4 reported here (Fig 9B, 9C). L797, G818, W821, H1838, and R1843 in ABCA4 are also present in ABCA1 and ABCA7 and are part of highly conserved motifs supporting the importance of these structural features in the proper folding and function of these transporters (Fig 9B,C, Fig S1A-B). In the case of H1838, four variants (H1838D, H1838N, H1838R, H1838Y) have been reported to cause STGD1 and shown here to have severe to moderate effects on ABCA4 functional properties suggesting that the histidine plays a crucial role in maintaining the optimal structure and function of ABCA4.

**Figure 9.**
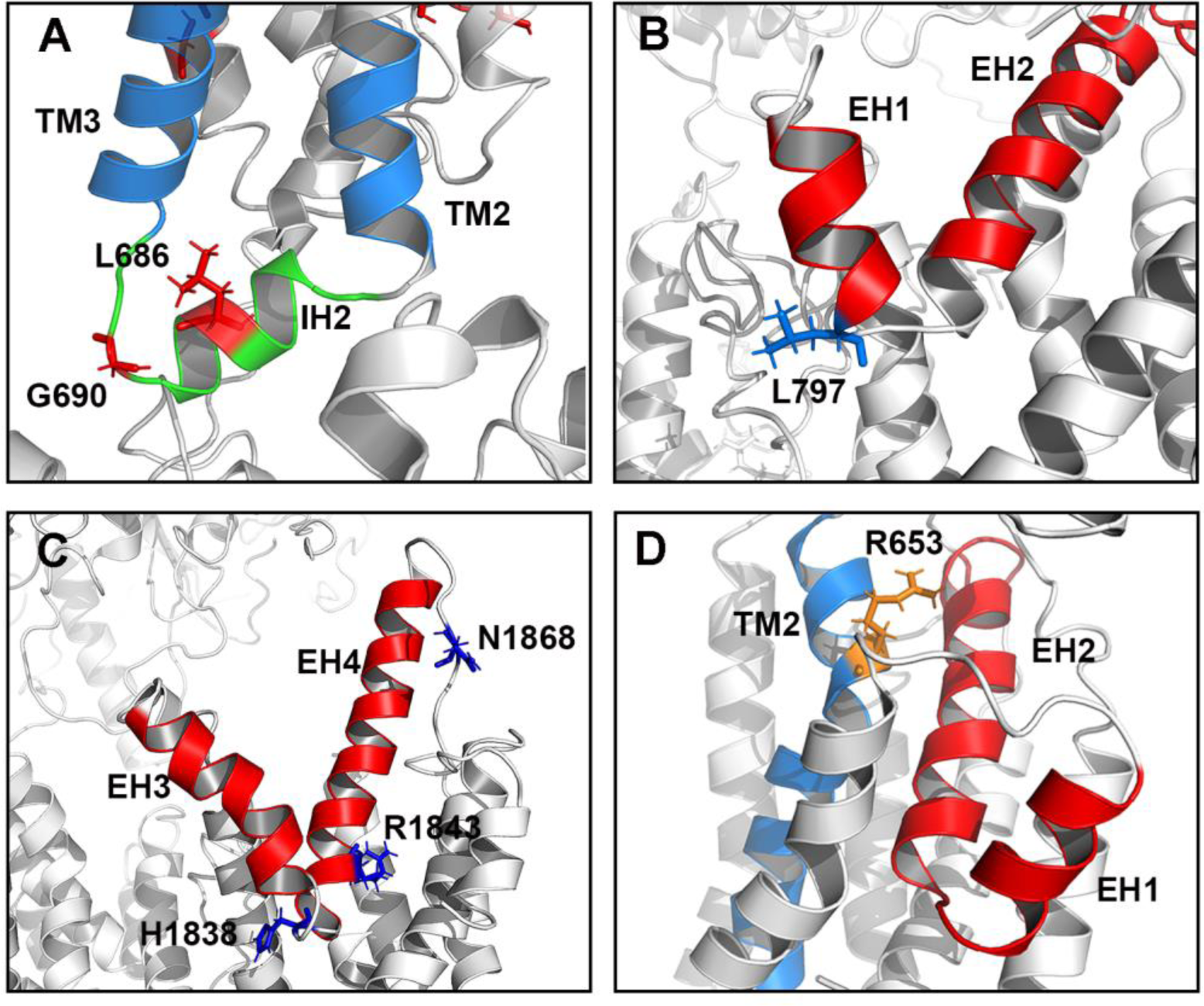
Location of selected amino acid residues (sticks) in the ABCA4 I-Tasser homology model that when mutated cause STGD1. **A**. The loop region between TM2 and TM3 (blue) showing the intracellular transverse helix IH2 (green) with residues L686 and G690. ABCA4 variants L686S and G690V cause severe protein misfolding and are predicted to cause severe STGD1 in trans with a null ABCA4 allele. **B**. The EH1 and EH2 helices (red) between TM5 and TM6 within TMD1 and showing the location of the L797 residue. The ABCA4 variant L797P causes significant protein misfolding and is predicted to cause severe STGD1 in trans with a null ABCA4 allele. **C**. The EH3 and EH4 helices (red) between TM11 and TM12 within TMD2 and showing the H1838, R1843 and N1868 residues. ABCA4 variants H1838D, H1838N, H1838R, and H1838Y are predicted to cause moderate or severe (see Table 2) STGD1 in trans with a null ABCA4 allele. ABCA4 variant R1843W is predicted to cause a moderate STGD1 in trans with a null ABCA4 allele. ABCA4 variant N1868I in the loop between EH4 and TM12 is predicted to cause a mild STGD1 phenotype based on functional assays. **D**. Location of R653 within TM2. This residue is suggested to comprise part of the binding site for *N*-Ret-PE substrate based on biochemical studies. TM2 (blue); EH1 and EH2 (red). R653C and R653H are predicted to cause severe and moderate STGD1, respectively.

Our classification based on functional data can be used to predict the impact that missense mutations have on the severity of STGD1. Unfortunately, sufficient metadata for large cohorts of patients including age of onset, visual acuity, autofluorescence, or other diagnostic properties have not been reported for most of the TMD mutations examined here to make reliable correlations between genotype and residual functional activity of the variants. However, such data are available in a few cases as for example P1380L Class 2 variant within TM7. The age of onset for individuals homozygous for this variant ranges from 10 to 26 years with an average age of 19 yr (Fingert, Eliason et al. 2006, Cideciyan, Swider et al. 2009, Hwang, Zernant et al. 2009, Tanna, Strauss et al. 2017). This age of onset is consistent with the moderate disease classification presented here and the moderate classification based on clinical findings (Fakin, Robson et al. 2016). When this mutation is in *trans* with the S1696N mutation, reported here to be a mild Class 3 mutation, then the age of onset is increased to 45 yr supporting our Class 3 designation of the S1696N variant (Hwang, Zernant et al. 2009).

N1868I is another mild variant for which there is considerable genetic and clinical data (Zernant, Lee et al. 2017, Runhart, Sangermano et al. 2018). The genetic frequency and penetrance of the N1868I mutations has led to considerable discussion about whether this variant is benign or pathogenic (Cremers, Lee et al. 2020). Support for the pathogenic nature of this variant comes from analysis of STGD1 patients harboring the N1868I variant in *trans* with a severe variant. The age of onset varies significantly between 18 to 72 even in patients with the same genotype (e.g. N1868I/L257Vfs*17)(Zernant, Lee et al. 2017, Runhart, Sangermano et al. 2018). Our biochemical studies indicate that the N1868I exhibits an expression and activity profile broadly similar to WT ABCA4. However, the affinity of this variant for *N*-Ret-PE is an order of magnitude lower than that of WT ABCA4 suggesting that this mutation alters the substrate binding properties of ABCA4. The relatively low affinity of this variant for *N*-Ret-PE would compromise the efficient transport of *N*-Ret-PE across disc membranes leading to some formation of bisretinoids in photoreceptors and gradual accumulation in RPE cells particularly when this variant is paired with a functionally deficient variant. The N1868I mutation is situated on a loop linking the V-shaped hairpin helices to TM12 in TMD2 (Fig 9C). Other mild variants include T716M, C764Y, V849A, E1399K, S1696N and R1898H/C. In some cases, a conserved amino acid substitution occurs as for example V849A, S1696N and R1898H, whereas in other cases the amino acid substitutions may have little impact on the structure or function of ABCA4 based on their location within the structure such as the E1399K or R1898C mutation.

Of the variants examined in this study, the R653C Class 1 variant is particularly interesting in that it shows unique properties. This variant solubilized in CHAPS buffer and localized within vesicle-like structures in transfected cells similar to WT ABCA4. Furthermore, the basal ATPase activity is similar to WT ABCA4. However, the *N*-Ret-PE binding and stimulated ATPase activity of this variant are severely impaired. These results strongly suggest that this mutation does not affect protein folding, but instead alters the interaction of *N*-Ret-PE with ABCA4. Additional substitutions at this position further highlight the requirement for an amino acid with a positively charged side chain at position 653 to facilitate *N*-Ret-PE binding. The R653A shows drastically reduced *N*-Ret-PE binding and ATPase activation and the R653E lacks these properties, whereas the R653K and R653H variants show significant substrate binding and ATPase activation. Although the R653H variant has significant substrate stimulated ATPase activity, this mutation has been reported to cause STGD1 (Cornelis, Bax et al. 2017). This may result from the lower affinity of this variant for *N*-Ret-PE (apparent K_d_ of 13.3µM for R653H vs 1.2µM for WT ABCA4) as shown in our binding studies. Our data suggest that the R653C with limited function would cause a more severe phenotype than R653H. This is supported by studies reporting that STGD1 patients compound heterozygous for the R653C and a frameshift mutation (T1537Nfs*18) or a premature stop mutation (R2030*) have an age of disease onset within the first decade of life (6-10 yrs old) (Huang, Zhang et al. 2013, Fujinami, Zernant et al. 2015). On the basis of these studies we suggest that arginine at position 653 located in TM2 close to the exocytoplasmic side of the membrane may form part of the binding pocket for *N*-Ret-PE. Modeling studies suggest that this binding site close to the interface with ECD1 may also comprise the helical segment (EH2) contained within the V-shaped hairpin structure between TM 5-6 TM5 (Fig 9D). In this respect it is interesting to note that in the structure of ABCA1, electron density that may represent the phospholipid substrate was observed within a shallow pocket enclosed by intracellular regions of TM1/2/5 (Qian, Zhao et al. 2017). The presence of the possible lipid binding site of ABCA1 at the interface with the cytoplasmic side and the proposed lipid binding site of ABCA4 on the exocytoplasmic side of the membrane is consistent with functional studies showing that ABCA1 is a lipid exporter with lateral access of the lipid substrate toward the cytoplasmic side and ABCA4 is a lipid importer with lateral access of the lipid substrate toward to exocytoplasmic side of the membrane (Quazi and Molday 2013). Additional studies are needed to more fully delineate the contributions of various amino acid residues within ABCA4 to *N*-Ret-PE binding.

In summary we present here expression and functional data for a large set of disease-associated mutations within the transmembrane domains of ABCA4. From these data, we have developed a classification which can be used to correlate the residual activity of the missense mutations with the severity of the disease. The information related to the functional properties and molecular mechanisms underlying TMD mutations should serve as a basis for the development of novel gene, drug and cell based therapies for STGD1 disease.

## Materials and Methods

### Prediction of Transmembrane Helices and Homology Model of ABCA4

The amino acid sequence of the transmembrane (TM) helices that form the TMDs of ABCA4 were predicted using algorithms based on hidden Markov models (DAS-TMfilter, ExPASy TMpred, HMMTOP, MP Toppred, PredictProtein, and TMHMM) (https://www.expasy.org/tools/). The TM predictions from each program were pooled together and the overlapping predictions common to all programs were used as the most probable TM sequences in the TMDs. To validate the quality of these predictions, we used TMD sequences reported for the cryo-electron microscopic structure of ABCA1 (Qian, Zhao et al. 2017).

Homology models of ABCA4 were made with I-TASSER and SWISS-MODEL using the ABCA1 Cryo-EM structure (5XYJ) as template, which bears 50% sequence identity with ABCA4 (Qian, Zhao et al. 2017). For modeling with I-TASSER, ABCA4 was split into two tandem halves and each half was modeled individually (Yang, Yan et al. 2015). The two halves had a 9 amino acid region of overlap just after the NBD1 (GDRIAIIAQ), and the two models were aligned and fused at this region of overlap using PyMol to construct the full homology model of ABCA4. Both programs gave comparable models.

### Cloning of ABCA4 Transmembrane Variants

The cDNA of human ABCA4 (NM_000350) containing a 1D4 tag (TETSQVAPA) at the C-terminus was cloned into the pCEP4 vector using the Nhe-I and Not-I restriction sites as previously described (Zhong, Molday et al. 2009). Missense mutations were generated by PCR based site-directed mutagenesis. All DNA constructs were verified by Sanger sequencing.

### Antibodies

The Rho 1D4 monoclonal antibody was generated in house (Molday and MacKenzie 1983) and used in the form of hybridoma culture fluid for Western blotting and immunofluorescence microscopy. For immunoaffinity purification, the purified Rho 1D4 antibody was covalently coupled to CNBr-activated Sepharose 2B as previously described (Zhong, Molday et al. 2009).

### Heterologous Expression of ABCA4 Variants in HEK293T Cells

HEK293T cells grown at 80% confluency in 10 cm plates were transfected with 10 µg of pCEP4-ABCA4-1D4 using 1 mg/ml PolyJet^™^ (SignaGen, Rockville, MD) at a 3:1 PolyJet^™^ to DNA ratio. After 6 to 8 h, the cells were placed in fresh media. At 48 h post-transfection, the cells were harvested and centrifuged at 2800 x g for 10 min. The pellet was resuspended in 200 µl of resuspension buffer (50 mM HEPES, 100 mM NaCl, 6 mM MgCl_2_, 10% glycerol, pH 7.4). A 40 µl aliquot of the resuspended pellet was solubilized at 4^°^C for 40 min in 500 µl of either 3-[(3-cholamidopropyl) dimethylammonio]-1-propanesulfonate hydrate (CHAPS) solubilization buffer (20 mM CHAPS, 50 mM HEPES, 100 mM NaCl, 6 mM MgCl_2_, 1 mM dithiothreitol (DTT), 1X ProteaseArrest, 10% glycerol, 0.19 mg/mL brain-polar-lipid (BPL), and 0.033 mg/mL 1,2-dioleoyl-sn-glycero-3-phospho-L-ethanolamine (DOPE) [Avanti Polar Lipids, Alabaster, AL], pH 7.4) or SDS solubilization buffer (3% SDS, 50 mM HEPES, 100 mM NaCl, 6 mM MgCl_2_, 1 mM DTT, 1X ProteaseArrest, 10% glycerol, 0.19 mg/mL BPL, and 0.033 mg/mL DOPE, pH 7.4). The samples were then centrifuged at 100,000 x g for 10 min in a TLA110.4 rotor using a Beckman Optima TL centrifuge. The supernatant was collected and the absorbance at 280 nm was measured to determine protein concentration. Approximately, 7-8 µg of total protein per lane were resolved on an 8% SDS-polyacrylamide gel and transferred onto a polyvinylidene difluoride membrane for Western blotting. The blots were blocked in 1% milk for 1 h and labelled for 2 h with culture fluid containing the Rho1D4 mouse monoclonal antibody (1:100 dilution in phosphate-buffer-saline (PBS)) and rabbit-anti-β-tubulin (1:1000 dilution in PBS) used as a loading control. The blots were washed 3 times with PBS followed by incubation for 1 h with donkey anti-mouse IgG or donkey anti-rabbit IgG conjugated to IR dye 680 (1:20,000 dilution in PBS-0.5%Tween (PBS-T). The blots were washed 3 times with PBS-T and imaged on an Odyssey Li-Cor imager (Li-Cor, Lincoln, NE). Protein expression levels were quantified from the intensity of the ABCA4 bands and normalized to the intensity of the bands from the β-tubulin loading control.

### Immunofluorescence Microscopy of ABCA4 Variants in Transfected COS-7 Cells

COS-7 cells were seeded 24 h before transfection on six-well plates containing coverslips coated with poly-L-lysine to promote cell adhesion to coverslips. The cells were transfected with 1 µg of DNA and 3 µl of 1mg/ml PolyJet^™^ for 6 to 8 h before replacing with fresh media. At 48 h post-transfection, the cells were fixed with 4% paraformaldehyde in 0.1M phosphate buffer (PB), pH 7.4, for 25 min and washed 3 times with PBS. The cells were then blocked with 10% goat serum in 0.2% Triton X-100 and PB for 30 min. Primary antibody labeling was carried out for 2 h in 2.5% goat serum, 0.1% Triton X-100 and PB using the Rho1D4 antibody against the 1D4-tag and the calnexin rabbit-polyclonal antibody as an endoplasmic reticulum (ER) marker. The coverslips were washed 3 times with PB, followed by secondary labeling using Alexa-488 goat-anti-mouse Ig (for ABCA4), Alexa-594 goat-antirabbit Ig (for calnexin), and counterstained for nuclei with DAPI for 1 h. The coverslips were washed 3 times with PB to remove excess antibody and subsequently mounted onto microscope slides with Mowiol mounting medium and kept in the dark at 4°C. The microscope slides were visualized under a Zeiss (Oberkochen, Germany) LSM700 confocal microscope using a 40X objective (aperture of 1.3). Images were analyzed using Zeiss Zen software and ImageJ.

### ATPase Assay

Depending on the ABCA4 variant, 1 to 3 ten cm plates of HEK293T at 80% confluency were transfected as described above. At 24 h post-transfection, the cells were harvested and centrifuged at 2800 x g for 10 min. Depending on the number of plates transfected, the pellet was resuspended in 1-2 ml of CHAPS solubilization buffer for 60 min at 4°C followed by a 100,000 x g centrifugation for 10 min as described above. The supernatant was incubated with 70 µl of packed Rho1D4-Sepharose affinity matrix for 60 min at 4°C. The beads were washed twice with 500 µl of column buffer (10 mM CHAPS, 50 mM HEPES, 100 mM NaCl, 6 mM MgCl_2_, 1 mM DTT, 1X ProteaseArrest, 10% glycerol, 0.19 mg/ml BPL, and 0.033 mg/ml DOPE, pH 7.4), transferred to an Ultrafree-MC spin column, and washed another five times with 500 µl of column buffer. Bound ABCA4 was eluted from the Rho1D4 matrix twice with 75 µl of 0.5 mg/ml 1D4 peptide in column buffer at 18°C. When necessary, the absorbance at 280 nm was taken to calculate the protein concentration and the samples were diluted with column buffer to have equal protein concentration across all variants tested. Thirty microliters of purified ABCA4 was loaded onto an 8% acrylamide gel along with BSA standards to calculate the amount of ABCA4 protein in each sample.

ATPase assays were carried out using the ADP-GLO^™^ Max Assay kit (Promega, Madison, WI) according to the manufacturer’s guidelines. For each ABCA4 variant tested, 15 µl aliquots of purified ABCA4 (∼100 ng of protein per tube) was added to six microcentrifuge tubes. One microliter of 0.8 mM all-*trans* retinal in column buffer (or column buffer alone) was added to half of the samples (done in triplicate) to obtain a final concentration of 0 or 40 µM all-*trans* retinal. Each tube was incubated for 15 min at room temperature in the dark to allow all-*trans* retinal to react with PE and form the Schiff base adduct *N*-Ret-PE. Subsequently, 4 µl of a 1 mM ATP solution (in column buffer) was added and the samples were incubated at 37°C for 40 min. The final concentrations of all-*trans* retinal and ATP in each sample were 40 µM (or 0 µM) and 200 µM, respectively. For all ATPase assays, the ATPase deficient mutant ABCA4-MM, in which the lysine residues in the Walker A motif of nucleotide binding domain 1 (NBD1) and nucleotide binding domain 2 (NBD2) were substituted for methionine, was used to subtract non-specific ATPase activity. Typically, the MM values were less than 10% of the basal ATPase activity.

### *N*-Ret-PE Binding Assay

Tritiated all-*trans* retinal was prepared by the method of Garwin and Saari (Garwin and Saari 2000, Zhong and Molday 2010). [^3^H] all-*trans* retinal was mixed with unlabeled all-*trans* retinal to obtain a final concentration of 1 mM and a specific activity of 500 dpm/pmol. For a typical binding assay, depending on the ABCA4 variant, 2-5 ten cm plates of transfected HEK293T cells were harvested 48 h post-transfection and centrifuged for 10 min at 2800 x g as described above. The pellet was resuspended and solubilized in 3 ml of CHAPS solubilization buffer for 40 to 60 min at 4°C and subsequently centrifuged at 100,000 x g for 10 min to remove unsolublized material. The supernatant was collected and divided in half. Each half was incubated with 80 µL of packed Rho1D4-Sepharose affinity matrix equilibrated in column buffer and mixed by rotation for 60 min at 4°C. The affinity matrix was washed twice with 500 µl of column buffer and mixed with 250 µl of 10 µM [^3^H] all-*trans* retinal (500 dpm/pmol) in column buffer for 30 min at 4°C. The matrix was then washed 6 times with 500 µl of column buffer. One sample was incubated with 1 mM ATP and the other was incubated in the absence of ATP for 15 min at 4° C. The affinity matrices were washed 5 times with 500 µl of column buffer and transferred to an Ultrafree-MC (0.45 µm filter) spin column followed by another 5 washes of 500 µl of column buffer. Bound [^3^H] all-*trans* retinal was extracted with 500 µL of ice-cold ethanol with shaking at 500 rpm for 20 min at room temperature and counted in a liquid scintillation counter. Bound ABCA4 was eluted from the Rho1D4-bead matrix with 3% SDS in column buffer and resolved in an 8% SDS-polyacrylamide gel for analysis of protein levels by Western blotting. All washes and incubations with [^3^H] all-*trans* retinal were done in the dark. In some studies, the concentration of all-trans retinal was varied between 0 and 50 µM.

## Acknowledgements

The authors thank Laurie Molday for valuable discussions related to the ATPase activity assays. This work was supported by grants from the Canadian Institutes for Health Research (PJT 148649 and the National Institutes of Health EY002422.

## Competing interests

The authors declare no competing interests.

## Notes

### Competing Interest Statement

The authors have declared no competing interest.

