## Supplemental Figures for "Pathogenic Mechanisms Underlying Stargardt Macular Degeneration Linked to Mutations in the Transmembrane Domains of ABCA4"

Supplemental Material

A

B

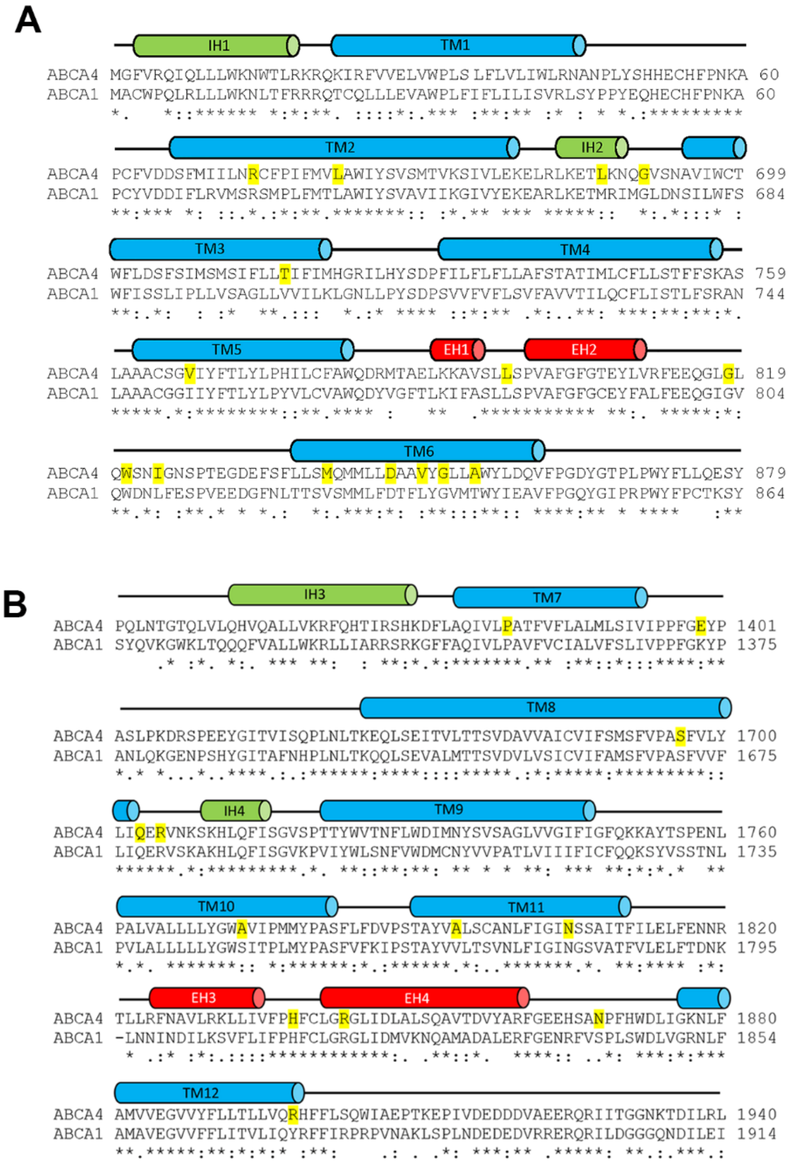


**Figure S1A-B: Sequence alignment of the Transmembrane Domains (TMDs) of ABCA4 and ABCA1 using Clustal Omega**. A. TMD1 showing the high degree of sequence similarity in the membrane spanning α-helical segments as blue cylinders (TM1-6), intracellular transverse coupling helices green (IH1 and IH2), and exoplasmic V-shaped α-helical hairpin helices in red (EH1 and EH2). B. TMD2 showing TM 7-12, IH3-4, and EH3-4. The disease-associated STGD1 missense mutations examined in this study are highlighted in yellow.

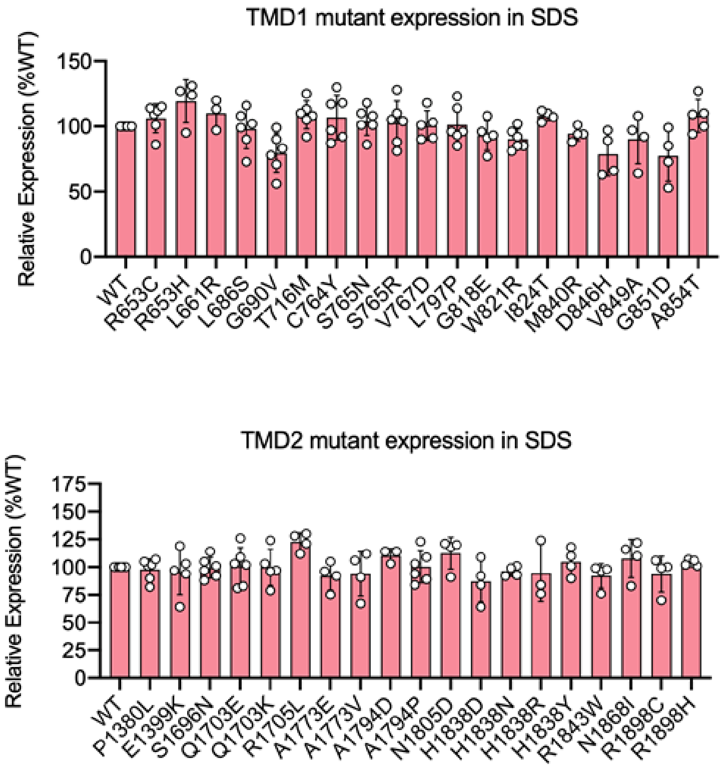


**Supplemental figure S2. Solubilization levels of TMD Variants in SDS buffer.** The solubilization levels of the TMD variants in SDS buffer were determined from Western blots of lysates from HEK293T cells expressing Wild-Type (WT) ABCA4 or variants with missense mutations in TMD1 (A) or TMD2 (B). Data is the mean ± SD for the indicated number of experiments. The majority of the variants expressed at within 80% of the WT level).
